# Intercellular adhesion stiffness moderates cell decoupling on stiff substrates

**DOI:** 10.1101/802520

**Authors:** D. A. Vargas, T. Heck, B. Smeets, H. Ramon, H. Parameswaran, H. Van Oosterwyck

## Abstract

The interplay between cell-cell and cell-substrate interactions is complex yet necessary for the formation and well-functioning of tissues. The same mechanosensing mechanisms used by the cell to sense its extracellular matrix, also play a role in intercellular interactions. We used the discrete element method to develop a computational model of a deformable cell that includes subcellular components responsible for mechanosensing. We modeled a cell pair in 3D on a patterned substrate, a simple laboratory setup to study intercellular interactions. We explicitly modeled focal adhesions between the cells and the substrate, and adherens junctions between cells. These mechanosensing adhesions matured; their disassembly rate was dictated by the force they carry. We also modeled stress fibers which bind the discrete adhesions and contract. The mechanosensing fibers strengthened upon stalling and exerted higher forces. Traction exerted on the substrate was used to generate maps displaying the magnitude of the tractions along the cell-substrate interface. Simulated traction maps are compared to experimental maps obtained via traction force microscopy. The model recreates the dependence on substrate stiffness of the tractions’ spatial distribution across the cell-substrate interface, the contractile moment of the cell pair, the intercellular force, and the number of focal adhesions. It also recreates the phenomenon of cell decoupling, in which cells exert forces separately when substrate stiffness increases. More importantly, the model provides viable molecular explanations for decoupling. It shows that the implemented mechanosensing mechanisms are responsible for competition between different fiber-adhesion configurations present in the cell pair. The point at which an increasing substrate stiffness becomes as high as that of the cell-cell interface is the tipping point at which configurations that favor cell-substrate adhesion dominate over those favoring cell-cell adhesion. This competition is responsible for decoupling. Additionally, we learn that extent of decoupling is modulated by adherens junction maturation.

**Statement of Significance:** Cells are sensitive to mechanical factors of their extracellular matrix while simultaneously in contact with other cells. This creates complex intercellular interactions that depend on substrate stiffness and play a role in processes such as development and diseases like cardiac arrhythmia, asthma, and cancer. The simplest cell collective system in vitro is a cell pair on a patterned substrate. We developed a computational model of this system which explains the role of molecular adhesions and contractile fibers in the dynamics of cell-cell interactions on substrates with different stiffness. It is one of the first models of a deformable cell collective based on mechanical principles. It recreates cellular decoupling, a phenomenon in which cells exert forces separately, when substrate stiffness increases.

## Introduction

Cells communicate with their environment and neighboring cells, transmitting forces through both. The interplay between cell-cell and cell-ECM adhesions is complex, with junctions sharing component proteins and being connected to one another via the cytoskeleton (1). Knowledge of this interplay is relevant to understand cell-cell communication in processes such as development and pathologies, including cardiac arrhythmia (2), asthma (3, 4), and cancer metastasis (5). Cell-pair studies on patterned substrates provide one of the simplest collective cell in vitro models. As previously presented in Polio *et al.*, in a pair of Airway Smooth Muscle (ASM) cells, cell-cell coupling strength decreases with increasing ECM stiffness while the total force exerted at the cell-cell interface via adhesions increases (6). Additionally, cell-cell junctions were visualized via immunostaining of β-catenin, a molecule in adherens junctions (AJs) cross-linking cadherins to the actin cytoskeleton. Visualization of β-catenin revealed that with increasing substrate stiffness, cell-cell AJs are progressively replaced by cell-ECM adhesions, such that the cell pair transitions from exerting force on the substrate as a single dipole into two separate ones (i.e. decouple) (6). A similar increase in cell-cell forces and replacement of cell-cell with cell-ECM adhesions was presented by McCain *et al.* for a cell-pair setup with cardiac myocytes (7). These findings raise the question: How does raising substrate stiffness on neighboring cells lead to the cells becoming individually stronger and exerting higher force at cell-cell adhesions, themselves stabilized by force, yet cause the cells to act separately on their substrate?

This decoupling has been observed not only for cell pairs, but also at a larger scale in endothelial sheets. Krishan *et al.* looked at gap formation in human umbilical vein endothelial cells (HUVECs) cultured on circular collagen islands (8). In these experiments, groups of 6-15 cells were cultured on substrates of different stiffness values in the range 1.2-90 kPa. As in cell-pair experiments, increased substrate stiffness led to higher cell-ECM and cell-cell forces. Staining of VE-cadherin and vinculin at cell-cell boundaries showed that the increased contractile forces, exerted on stiffer substrates, did not imply higher expression of either VE-cadherin or vinculin. Enhanced formation of F-actin stress fibers via thrombin treatment, however, did cause enhanced expression of vinculin (appearing in a dotted pattern at the cell-cell boundaries) and gap formation between cells. Thrombin is an inflammatory mediator which causes increased actomyosin contractility (9).

In another study, by Kugelmann *et al.*, actomyosin contractility was enhanced in human dermal microvascular endothelial cell (HDMEC) confluent sheets via use of either thrombin or histamine, and α-catenin and vinculin were imaged (10). Histamine, like thrombin, is a mediator involved in inflammatory responses and acts on myosin light chain kinase (MLCK) enhancing cellular contraction. α-catenin is a protein that binds β-catenin, together cross-linking VE-cadherin and F-actin; in this study, imaging is done via an anti-body which binds a particular epitope of α-catenin (i.e. alpha 18 subunit of catenin) that is exposed during tension. Thus an increase in α-catenin detection implies an increase in local force exertion (11). Mean changes of 201.1% ± 10.0% and 189.7 ± 9.2% in the intensity of α-catenin were measured 3 min after addition of histamine and 5 min after addition of thrombin, respectively. As in the work performed by Krishnan *et al.*, treatment with mediators resulted in vinculin becoming more visible in a dotted pattern at cell-cell boundaries. In both cases there was gap formation and increased stress fiber formation. In Kugelmann *et al.*, cellular decoupling was identified as an increase in transendothelial electric resistance of the monolayer. Thus both studies on endothelial sheets show that despite cell-cell adhesion reinforcement with increased actomyosin force, as evidenced by recruitment of additional molecules to the adhesion complex (i.e. vinculin or α-catenin), cells can become decoupled from one another.

To provide insight into the coexistence of cell decoupling and adhesion reinforcement, we developed a computational model of a cell-pair on a rectangular pattern in which subcellular force exertion mechanisms are taken into account, specifically: adhesion complex maturation (i.e. stabilization of adhesion with force carried) and stress fiber strengthening (i.e. increase in contraction force after stalling in fiber shortening). We compare simulated traction maps with those recovered in vitro through traction force microscopy (6). In the model both cell-ECM and cell-cell bonds are stabilized by force. The model captures the drop in cell coupling with increasing substrate stiffness and simultaneous increase in cell adhesion observed experimentally. We show that this occurs only over a specific range of substrate stiffness values with the stiffness of AJs near its center. The relative sensitivity to force of focal adhesions (FAs) and AJs does regulate this phenomenon. Altogether, our results draw a clearer picture of the effects of cellular mechanics in cell decoupling.

## Methods

The system studied via a computational model consists of two identical cells placed next to each other on a rigid substrate plane. The discrete element method (DEM) is used to represent both cells and the substrate via lattice-free nodes, connected to each other in such a way that each body (i.e. two cells and substrate plane) consists of a mesh of triangular elements where the nodes are the vertices of the triangles; triangles constituting the cells and substrate surfaces are used to calculate mechanical interactions between the different bodies in the system (12). The system is initialized such that the cells are spread on the surface and in contact with each other, thus cell-cell and cell-ECM interfaces exist at the start of the simulations. In this way the simulations represent the system observed experimentally in which cell pairs are in contact with one another on patterned ligand substrates (6). A visualization of the cell and substrate, displaying the meshing and substrate patterning can be found in the Supporting Material (Figure S3). An interval of 4h was simulated, long enough to ensure that a dynamic equilibrium was reached by the cell pair for analysis purposes. Figure 1 consists of a schematic of the model presenting the different elements of the cells and the forces involved in the system. Formulation and implementation are presented next. Model parameters are listed along with their values and source of estimation in Table 1.

**Table 1.**
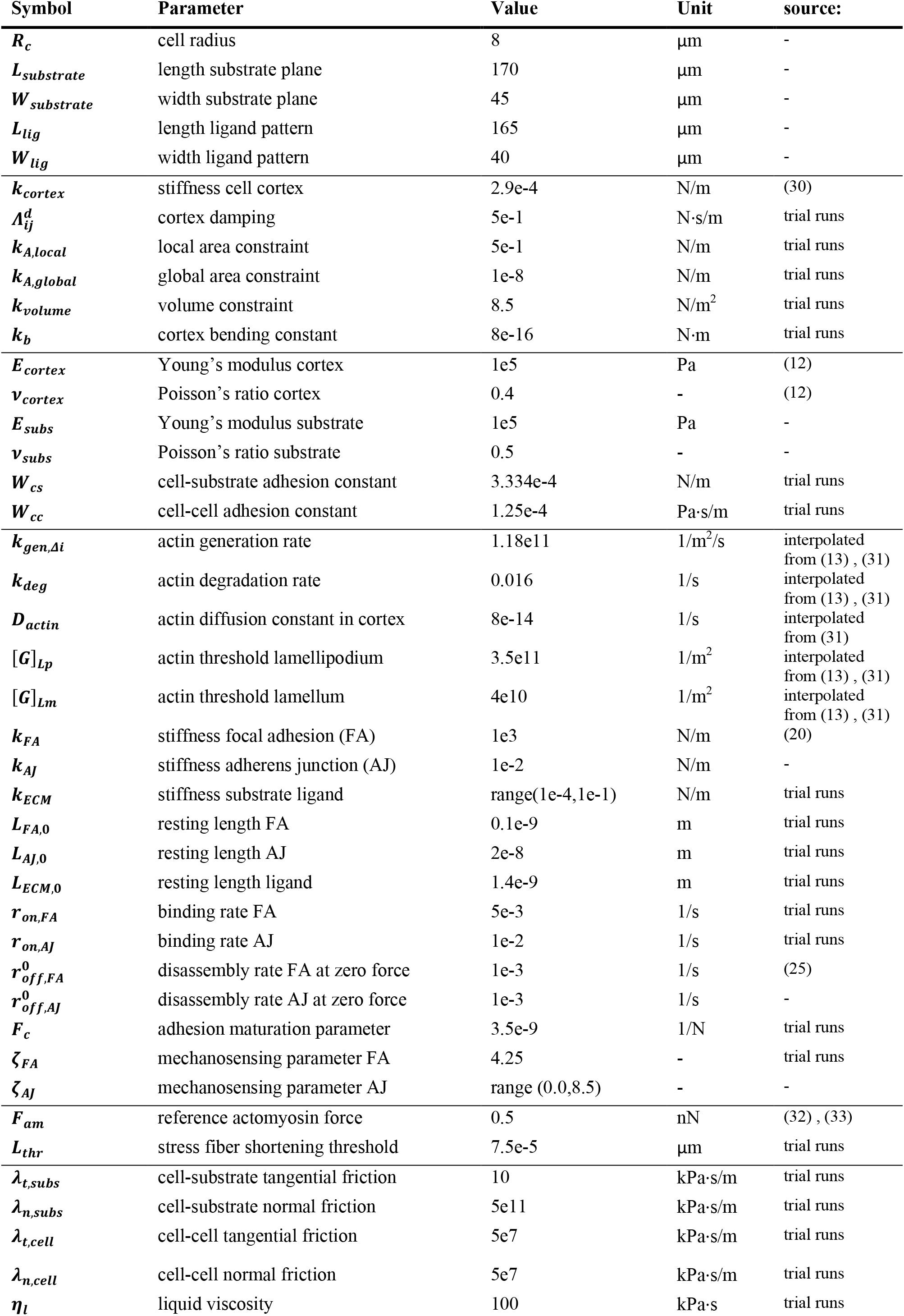
Model parameters

| Symbol | Parameter | Value | Unit | source: |
| --- | --- | --- | --- | --- |
| $R_c$ | cell radius | 8 | $\mu\text{m}$ | - |
| $L_{\text{substrate}}$ | length substrate plane | 170 | $\mu\text{m}$ | - |
| $W_{\text{substrate}}$ | width substrate plane | 45 | $\mu\text{m}$ | - |
| $L_{\text{lig}}$ | length ligand pattern | 165 | $\mu\text{m}$ | - |
| $W_{\text{lig}}$ | width ligand pattern | 40 | $\mu\text{m}$ | - |
| $k_{\text{cortex}}$ | stiffness cell cortex | 2.9e-4 | N/m | (30) |
| $\Lambda_{ij}^d$ | cortex damping | 5e-1 | N·s/m | trial runs |
| $k_{A,\text{local}}$ | local area constraint | 5e-1 | N/m | trial runs |
| $k_{A,\text{global}}$ | global area constraint | 1e-8 | N/m | trial runs |
| $k_{\text{volume}}$ | volume constraint | 8.5 | N/m<sup>2</sup> | trial runs |
| $k_b$ | cortex bending constant | 8e-16 | N·m | trial runs |
| $E_{\text{cortex}}$ | Young's modulus cortex | 1e5 | Pa | (12) |
| $\nu_{\text{cortex}}$ | Poisson's ratio cortex | 0.4 | - | (12) |
| $E_{\text{subs}}$ | Young's modulus substrate | 1e5 | Pa | - |
| $\nu_{\text{subs}}$ | Poisson's ratio substrate | 0.5 | - | - |
| $W_{cs}$ | cell-substrate adhesion constant | 3.334e-4 | N/m | trial runs |
| $W_{cc}$ | cell-cell adhesion constant | 1.25e-4 | Pa·s/m | trial runs |
| $k_{\text{gen},\Delta i}$ | actin generation rate | 1.18e11 | 1/m<sup>2</sup>/s | interpolated from (13) , (31) |
| $k_{\text{deg}}$ | actin degradation rate | 0.016 | 1/s | interpolated from (13) , (31) |
| $D_{\text{actin}}$ | actin diffusion constant in cortex | 8e-14 | 1/s | interpolated from (31) |
| $[G]_{Lp}$ | actin threshold lamellipodium | 3.5e11 | 1/m<sup>2</sup> | interpolated from (13) , (31) |
| $[G]_{Lm}$ | actin threshold lamellum | 4e10 | 1/m<sup>2</sup> | interpolated from (13) , (31) |
| $k_{FA}$ | stiffness focal adhesion (FA) | 1e3 | N/m | (20) |
| $k_{AJ}$ | stiffness adherens junction (AJ) | 1e-2 | N/m | - |
| $k_{ECM}$ | stiffness substrate ligand | range(1e-4,1e-1) | N/m | trial runs |
| $L_{FA,0}$ | resting length FA | 0.1e-9 | m | trial runs |
| $L_{AJ,0}$ | resting length AJ | 2e-8 | m | trial runs |
| $L_{ECM,0}$ | resting length ligand | 1.4e-9 | m | trial runs |
| $r_{on,FA}$ | binding rate FA | 5e-3 | 1/s | trial runs |
| $r_{on,AJ}$ | binding rate AJ | 1e-2 | 1/s | trial runs |
| $r_{off,FA}^0$ | disassembly rate FA at zero force | 1e-3 | 1/s | (25) |
| $r_{off,AJ}^0$ | disassembly rate AJ at zero force | 1e-3 | 1/s | - |
| $F_c$ | adhesion maturation parameter | 3.5e-9 | 1/N | trial runs |
| $\zeta_{FA}$ | mechanosensing parameter FA | 4.25 | - | trial runs |
| $\zeta_{AJ}$ | mechanosensing parameter AJ | range (0.0,8.5) | - | - |
| $F_{am}$ | reference actomyosin force | 0.5 | nN | (32) , (33) |
| $L_{thr}$ | stress fiber shortening threshold | 7.5e-5 | $\mu\text{m}$ | trial runs |
| $\lambda_{t,subs}$ | cell-substrate tangential friction | 10 | kPa·s/m | trial runs |
| $\lambda_{n,subs}$ | cell-substrate normal friction | 5e11 | kPa·s/m | trial runs |
| $\lambda_{t,cell}$ | cell-cell tangential friction | 5e7 | kPa·s/m | trial runs |
| $\lambda_{n,cell}$ | cell-cell normal friction | 5e7 | kPa·s/m | trial runs |
| $\eta_l$ | liquid viscosity | 100 | kPa·s | trial runs |

**Figure 1.**
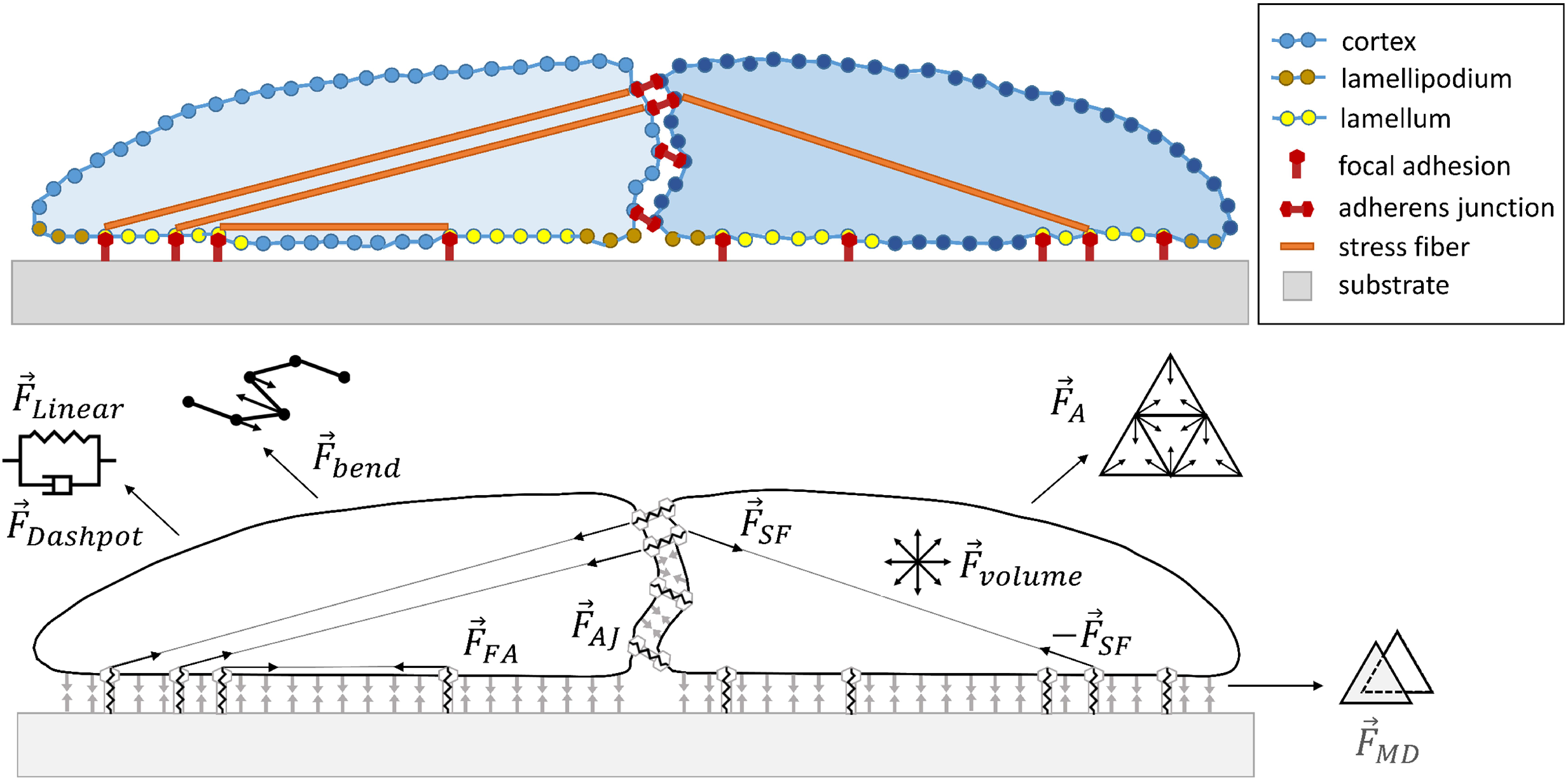
Schematic of a cross-section of cell pair displaying the different parts represented in the model (top). Forces involved in evolution of the mechanical system (bottom). The following relevant forces are indicated: Cortical elastic spring 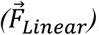, cortical dissipation 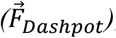, cortical bending 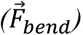, local triangle and global cell area conservation 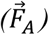, cell volume conservation 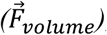, membrane contact 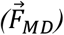, focal adhesion 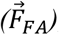, adherens junction 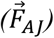, and stress fiber 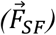.

### Cell anatomy

Triangles are also used to demarcate distinct parts of the cellular anatomy that exist at the cell-substrate interface, specifically a lamellipodium and a lamellum. The former is the outer most area, where a protrusion force would be exerted outwards toward the cell perimeter by polymerization of actin (driving retrograde flow) and pushing off nascent adhesions in cell migration. The lamellipodium has a width that varies between 1-3 μm. Where the lamellipodium ends, the lamellum starts, also radially located in the cell-substrate interface. In the lamellum, actin flow slows down allowing for formation of stress fibers biding mature FAs; the lamellum has a thickness of around ~12 μm (13). A representation of the lamella of the cells in the cell pair can be found in the Supporting Material (Figure S3). Beyond these two areas, the inner part of the cell-substrate interface is a part of the cell body in which no protrusion or formation of FAs can occur.

To demarcate these areas of the cell-substrate interface, the cellular perimeter is first identified. Triangles in the perimeter act as sources of actin. Actin diffuses across all other triangles with a diffusion constant (*D*_*actin*_) in an act representative of actin retrograde flow. Together with diffusion, the concentration of actin at each triangle *i* (*G*_*Δi*_) (units of molecules/μm^2^) is determined by generation at perimeter) and degradation of actin; all triangles act as sinks for actin. This effectively creates a gradient of actin concentration that is highest at the edge and decays along the bottom and top surfaces of the cell. The change in concentration of actin at each triangle is given by Equation 3:

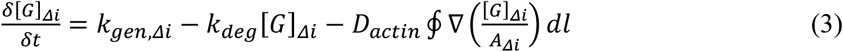

where *k*_*gen*,Δ*i*_ is the rate at which actin is generated per surface area unit at the edge triangles of the cell, and *k*_*deg*_ the rate at which actin is degraded in all triangles. The third term in the right-hand side of the equation corresponds to the change in concentration of actin at triangle *i* with area *A*_*Δi*_ due to diffusion across the sides of the triangle *l*. By setting two threshold values the lamellipodium and lamellum areas are dynamically defined in simulations, [*G*]_*Lp*_ and [*G*]_*Lm*_: At every time step the concentration of actin at each triangle *i* is checked, if [*G*]_*Δi*_ > [*G*]_*Lp*_ then the triangle is part of the lamellipodium, and if [*G*]_*Lp*_ > [*G*]_*Δi*_ > [*G*]_*Lm*_ then the triangle is part of the lamellum. If [*G*]_*Δi*_ < [*G*]_*Lp*_ the triangle is considered part of the cell body.

### Deformable cell model

Each of the cells is generated by subdividing an icosahedron and projecting the nodes onto a sphere (14). Using five subdivisions, this corresponds to a total of 2562 nodes (5120 triangles) for a cell. The radius of the cell as a sphere, before spreading, is *R*_*c*_ = 8 μm. The mesh represents the actin cortex of the cell underlying the cell membrane as the connections between nodes are made viscoelastic by connecting the nodes via Kelvin-Voigt elements (i.e. an elastic spring and viscous damper in parallel). A linear elastic spring is used, with Equation 1 describing the magnitude of the force acting on the vertices connected by a line element:

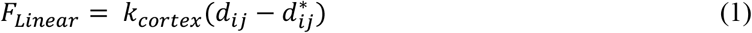

where *d*_*ij*_ and 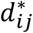 are the actual distance and equilibrium distance between nodes *i* and *j* (vertices), and *k*_*cortex*_ is the spring constant of the cellular cortex. The magnitude of the force contributed by the dashpot is described by Equation 2:

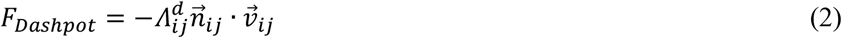

where 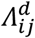 is the damping constant and 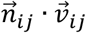 the projection of the velocity along the connecting axis between nodes *i* and *j*. Additional forces defining mesh geometry include a local triangle and global cell area conservation (*F*_*A*_), a cell volume conservation (*F*_*volume*_), and a resistance to bending based on the angle between the two planes defined by neighboring triangles (*F*_*bend*_). The details on implementation of these forces describing the cortex elasticity can be found in the Supporting Material.

### Substrate plane

The substrate is modeled as a rigid plane. It is a rectangle with dimensions of 170 × 45 μm^2^ (*L*_*substrate*_ × *W*_*substrate*_); it is also triangulated, using right isosceles triangles with area of 2.102 μm^2^. Triangulation is used to define smaller areas to which the cells can attach, representative of ligand patterning. For this study, the pattern was a smaller concentric rectangle of 165 × 40 μm^2^ (*L*_*lig*_ × *W*_*lig*_). This was done to avoid that the cells attached all the way to the end of the substrate plane but still limit the area to where the cells could attach. Substrate triangles are also used to discretize the area of the substrate and calculate the area of interaction between cell and substrate triangles, as well as record local forces applied on the substrate to generate a traction map that shows the spatial distribution of tractions exerted on the substrate by the cell pair.

### Cell adhesion

Two types of interactions between cells and between cells and substrate are modeled: transient and multi-protein complexes. The former represents transient binding of integrin molecules on the cell surface with ligands on the substrate (in the case of cell-substrate adhesion) and cadherin molecules on the membrane of different cells (in the case of cell-cell adhesion). The latter represents discrete mature molecular adhesions that have matured into multi-protein complexes: FAs (in the case of cell-substrate adhesion) and AJs (in the case of cell-cell adhesion).

Transient adhesion between triangles that are in contact is modeled according to Maugis-Dugdale theory This theory describes interactions based on spherical overlap, so interaction force 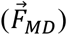 between a surface and its surroundings is computed as the interactions between two spheres corresponding to two triangles in contact. Maugis-Dugdale theory includes both a Hertzian elastic interaction (repulsion) for a radius of interaction *a* (based on sphere overlap), and an extended radius *b* (where *b* > *a*) in which an attraction potential is defined. To do this the local curvature of the triangulated surface is calculated based on local node coordinates; this curvature is then used to fit a unique sphere to each triangle. Spheres have a Young’s modulus (*E*_*cortex*_ and *E*_*subs*_ for cell and substrate respectively) and Poisson’s ratio (*v*_*cortex*_ and *v*_*subs*_ for cell and substrate respectively). The attraction constant describing the attraction potential is defined for cell-substrate (*W*_*cS*_) and cell-cell (*W*_*cc*_) transient interactions. An in-depth description of how Maugis-Dugdale theory has been applied in the context of contact mechanics can be found in the Supporting Material. This force, applied at triangle-triangle contacts and accounting for both attraction and repulsion, ensures that cells spread on the substrate plane and that the cell-cell interface remains smooth.

Multi-protein complexes are modeled through discrete elements at the cell nodes of the triangular mesh. In the case of cell-substrate interaction, a node in the lamellum area can form a discrete adhesion (representative of a FA) with any point on the rectangular pattern (representative of ligand on the substrate plane) as long as it is within a (vertical) distance of 0.075 μm to the plane. In the case of cell-cell interaction, any two nodes in different cells within 0.75 μm of each other can form a discrete adhesion (representative of an AJ). This maximum interaction distance is larger for cell-cell interactions to ensure that likelihood of forming each type of adhesion is similarly dependent on a binding rate, since a node binding the substrate can bind anywhere on the plane (independent of meshing) while a node in proximity of another cell can only bind to another node.

Discrete adhesions, representative of multi-protein complexes, are implemented as non-compressible Hookean springs. In the case of FAs, a two-spring system (in series) is used, a representation already proposed by Schwarz *et al.* (16): A stiff spring represents the FA (spring constant *k*_*FA*_), while a softer spring represents the underlying ligand molecule (spring constant *k*_*ECM*_). The force carried by the two spring system describing the FA at node *i* 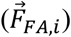 is described by Equation 4:

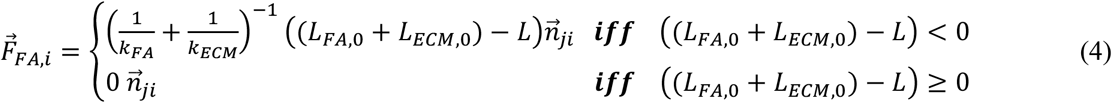

where *L*_*FA*,0_ and *L*_*ECM*,0_ are the equilibrium lengths of the FA and ECM ligand fiber, and *L* is the length at each corresponding time step. 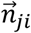 is the unit vector in the axis that runs from point *j* in the substrate (not necessarily a node) to cell node *i*. As *k*_*FA*_ ≫ *k*_*ECM*_, the FAs force response is dictated by *k*_*ECM*_ (see Table 1). The spring stiffness for the ligand can be converted to a bulk stiffness felt locally by the cell value according to Equation 5, taken from Mitrossilis *et al.* (17):

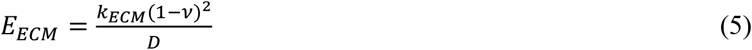

where *E*_*ECM*_ is the Young’s modulus of the substrate, *v* is its Poisson’s ratio, and *D* is the diameter of the contact area of the FA with the substrate. *v* was set to 0.5, and *D* was set to 1 μm (which approximates the variable diameter observed throughout simulations). *E*_*ECM*_ will be assumed to be the substrate stiffness the simulated cell senses to compare our output to experimental results, given the limitation of using a rigid plane as the substrate; This assumes that each FA only senses the substrate with which it is directly in contact. In this way different stiffness conditions can be modeled despite using a rigid plane as the substrate. The values of the spring constant used to define ECM stiffness were: *k*_*ECM*_ = [1e-4, 3e-4, 5e-4, 1e-3, 2e-3, 3e-3, 5e-3, 1e-2, 2e-2, 3e-2, 5e-2, 1e-1] N/m. This corresponds to a bulk stiffness values of: *E*_*ECM*_ = [0.025, 0.075, 0.125, 0.25, 0.5, 0.75, 1.25, 2.5, 5, 7.5, 12.5, 25] kPa.

In the case of AJs, a single non-compressible Hookean spring is used to describe the adhesion (spring constant *k*_*AJ*_). The force carried by a single adhesion at node *i* 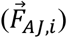 is described in Equation 6:

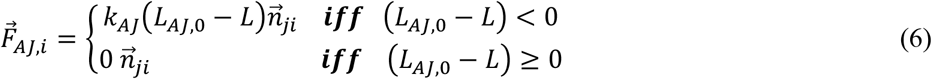

where *L*_*AJ*,0_ is the equilibrium length of the spring representing a AJ, and *L* is the length at each corresponding time step. 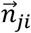 is the unit vector in the axis that runs from node *j* on one cell to node *i* on another cell. At each node, if in proximity of a binding target, a discrete adhesion can be formed based on a binding rate for FAs (*r*_*on*,*FA*_) and AJs (*r*_*on*,*AJ*_) in a stochastic fashion. Similarly a rate dictates how often adhesions are disassembled for FAs (*r*_*off*,*FA*,*i*_) and AJs (*r*_*off*,*AJ*,*i*_); however, this rate is made dependent on the magnitude of the force carried by the adhesion in such a way that force stabilizes the adhesion reducing the disassembly rate. This is the first way in which mechanosensing is accounted for in this model, and it is based on findings showing that multi-protein adhesion complexes are capable of mechanosensing and are stabilized by force through protein recruitment (11, 18, 19). Based on modeling of catch bonds (20), Equations 7 describe the relation between disassembly rate and force carried by the adhesions:

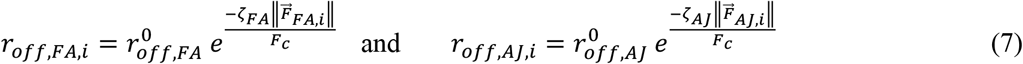

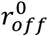 signifies the rate of disassembly when no force is carried by an adhesion; ζ and *F*_*c*_ are parameters that control the degree of mechanosensing. Figure 2A shows the effect of varying the parameter ζ_*AJ*_ on disassembly rate of AJs and corresponding expected adhesion lifetime values as a function of force. If a focal adhesion is outside of the lamellum (which can occur due to displacement of the cell nodes with force or change in shape of the lamellum) the force dependent rate of disassembly (*r*_*off*,*FA*,*i*_) is increased by a factor of 10.

**Figure 2.**
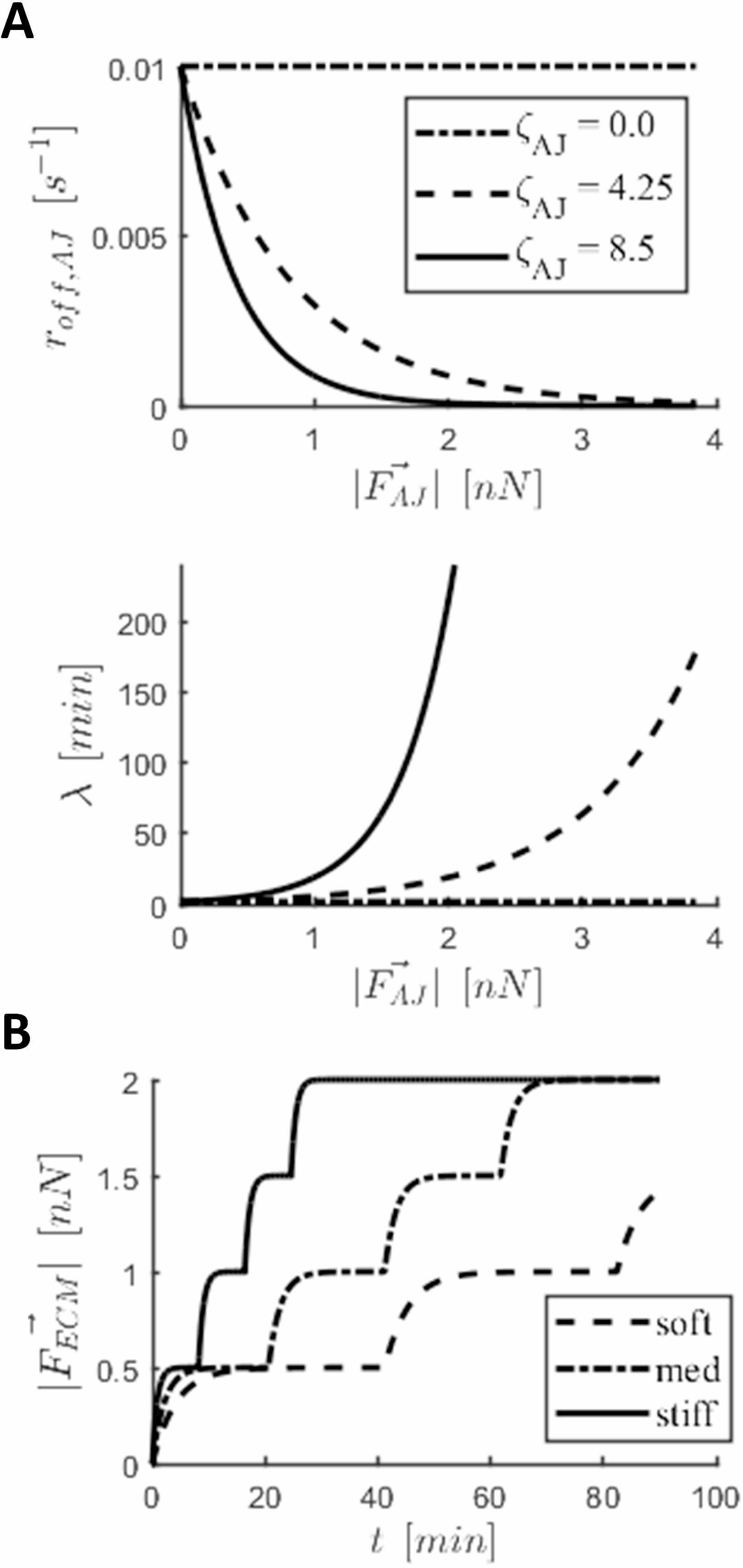
Implementation of mechanosensing mechanisms in the cell. A) Adherens junction disassembly rate (r_off,AJ_) decreases with force carried by adhesion 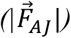(top). Corresponding expected adhesion lifetimes (λ), where r_off,AJ_ = 1 − e^−1/λ^ (bottom). Parameter ζ_AJ_ dictates sensitivity to force. B) Step increase in force carried by ECM 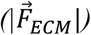 abound to a focal adhesion and applied by a stress fiber. Steps correspond to stalling in fiber contraction. The stiffer the substrate, the faster a fiber strengthens taking more steps during its lifetime. Graphs shown are a result of a simulation of an isolated two-spring system (ideal case).

### Stress fibers

Stress fibers are simulated as a pair of equal and opposite forces acting on a pair of discrete adhesions (either two FAs or a FA and an AJ) attracting them to one another. These forces represent tension carried by stress fibers arising from the action of myosin II motors sliding along antiparallel actin bundles constituting the fiber. A fiber will form within a cell, connecting two adhesions chosen at random, if the potential fiber meets three conditions:

1. Adhesion is not already bound to a fiber
2. The fiber has a minimum initial length of 11 μm
3. The orientation angle of the fiber is equal or lesser than π/10 rad: This angle is the intersection of the projection of the fiber on the substrate plane and the long side of the rectangular ligand pattern.

Conditions 2 and 3 were set based on observations in the literature that fibers will only form along the long axis of a rectangular underlying pattern (6, 7). A fiber remains bound to the nodes and exerts force as long as both nodes are engaged in a discrete adhesion; as soon as one of the two adhesions is disassembled, the fiber (and corresponding force) is deleted. A visualization of the location of FAs within the lamellum, AJs in the cell-cell interface, and stress fibers bound the these adhesions can be found in the Supporting Material (Figure S3).

When formed, fibers have a set force magnitude (*F*_*am*_, where *am* stands for actomyosin). The force exerted by the fiber brings the adhesions connected via the fiber closer to one another. The distance between adhesions (i.e. fiber length) is calculated at every time step and the values averaged over 10 second intervals; this is done to obtain a value of fiber length that is stable despite fluctuations due to all the different forces acting on each node. When the length ceases to change for a particular fiber *f*, a factor describing fiber strengthening (*n*_*str*,*f*_) is incremented. This stalling in fiber length corresponds effectively to the moment when the difference between subsequent (i.e. 10 s apart) averaged values of the length falls below a threshold (*L*_*thr*_). The force exerted by stress fiber *f* on node *i*, when connecting nodes *i* and *j*, is described in Equation 8:

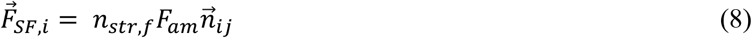

where 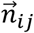 is the unit vector in the axis that runs from node *i* to node *j*. Upon formation, fibers have a value of *n*_*str*,*f*_ = 1. With each stalling event, this factor is increased by the value of 1, up to a maximum of 5 (under the assumption that that there is a limit to the force exerted by a stress fiber).

This strengthening of stress fibers is another mechanism of mechanosensing found in cells; it is based on the experimental findings by Wolfenson *et al.* (21) and a previous theoretical work by Parameswaran *et al.* (22). Wolfenson *et al.* showed that there is a stepwise strengthening of actomyosin fibrils, concurrent with recruitment of α-actinin molecules to FAs upon stalling of underlying substrate deformation.. The authors suggest that simultaneous recruitment of myosin to the fibrils is responsible for their strengthening. The interval between maturation steps depended on substrate stiffness, with strengthening occurring more often on stiffer substrates (22). These studies explain how cells exert higher tractions on stiffer substrates. An ideal dependence of fiber strengthening on substrate stiffness is shown in Figure 2B, which shows the evolution of the force carried by the two-spring system 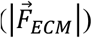 representing the FA and ligand (i.e. ECM) as the force exerted by a single stress fiber increases in steps triggered by fiber stalling. The experimental findings by Wolfenson *et al.* are recreated through the stress fiber strengthening mechanisms implemented in this model; fibers will stall faster in cells on stiffer substrates (rate of force increase is higher (16)) and thus undergo a strengthening step more often than fibers in cells on softer substrates. Thus, in the same time span, fibers will strengthen more and exert more force in cells on stiffer substrates.

### Equation of motion

Evolution of the system is described by the equation of motion. Because cells occupy a low Reynold’s (overdamped) environment where inertial forces are negligible (23), conservation of momentum for each node *i* takes the form of Equation 9:

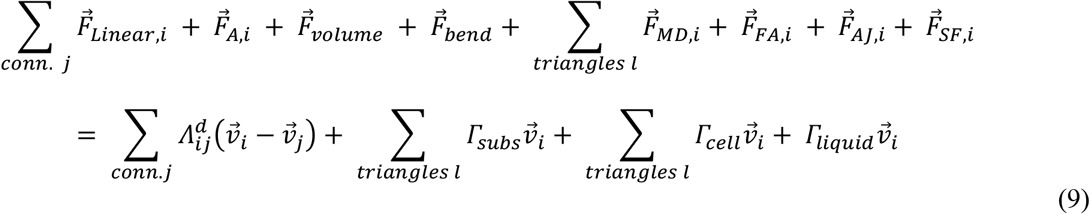

The left-hand side contains the sum of all forces acting on the node. The linear force due to the springs in the Kelvin-Voigt model are summed for each node over all connections (conn.). For the Maugis-Dugdale force, as it acts on the triangles in the mesh, forces are transfixed to the nodes (12).

The right-hand side of the equation shows the viscous friction, which in an overdamped environment is described by the product of friction acting on node *i* and its velocity 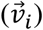. Different sources of friction include: Dissipation of the actin cortex by all dampers *j* connected to node *i* according to damping constant 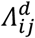; Friction due to contact with substrate triangles (*Γ*_*subs*_), a function of tangential and normal friction coefficients (*λ*_*t*,*subs*_ and *λ*_*n*,*subs*_, respectively); Friction due to contact with triangles of the other cell (*Γ*_*cell*_), a function of tangential and normal friction coefficients (*λ*_*t*,*cell*_ and *λ*_*n*,*cell*_, respectively); and Stokes’ drag (*Γ*_*liquid*_), a function of the liquid’s viscosity (*η*_*l*_). A mathematical description of the dissipative forces in terms of friction coefficients and medium’s viscosity can be found in the Supporting Material.

Equation 9 is a first order differential equation that couples the movements of all nodes. The system is solved iteratively for the velocities using the conjugate gradient method. More information on the numerical solution of the system can be found in the Supporting Material. A time step of 0.05 s was used in all simulations. All simulations are performed using the C++ particle-based software Mpacts (http://mpacts.com). Each simulation was run using an Intel Xeon Processor E5-2680 v3 on a node with 2.7 GB of memory; simulations were run in parallel utilizing the multiple nodes (20) per core. Each simulation took approximately 12 h to run.

## Results and Discussion

Each simulation consisted of a cell pair interacting with one another on a rectangular patterned substrate for a period of 4h. The stiffness of the substrate (*E*_*ECM*_) as well as the parameter controlling the degree of mechanosensing of AJs (ζ_AJ_) were varied to define the different conditions. The following values of substrate stiffness were used: *E*_*ECM*_ = [0.025, 0.075, 0.125, 0.25, 0.5, 0.75, 1.25, 2.5, 5, 7.5, 12.5, 25] kPa. To explore the effect of mechanosensing at the adhesion complexes on cell pair coupling and its dependence on substrate stiffness, we varied ζ_*AJ*_ relative to ζ_*FA*_. The difference in the values of ζ for each type of adhesion can represent the difference in the force sensitive molecules responsible for stabilization with force, α-catenin in AJs (11) and talin in FAs (24), for example. Three different scenarios were considered based on the relative difference in these parameters, such that we explore what occurs when AJs are not mechanosensing, when FAs and AJs are equally mechanosensing, and when AJs are relatively more mechanosensing. These scenarios and corresponding values of parameter ζ were: ζ_FA_(4.25) > ζ_AJ_(0.0), ζ_FA_(4.25) = ζ_AJ_(4.25), and ζ_FA_(4.25) < ζ_AJ_(8.5). ζ_FA_ was kept constant for easier comparison between scenarios, since we focus on cell-cell force transmission; also, mechanosensing of FAs has been further explored in the literature.

Qualitatively, decoupling with increasing ECM stiffness can be observed by the redistribution of traction forces along the cell-substrate interface. This is shown in Figure 3, where characteristic traction maps of cell pairs on substrates of varying stiffness along with cellular outlines are displayed: On soft substrates the cells are coupled and force is mostly exerted by stress fibers connecting FAs in the outer cellular lamella (i.e. farthest away from cell-cell interface) and AJs at the cell-cell interface. In the traction map, this means the highest tractions are located at the edges of the cell pair. As the substrate stiffness increases there is a progressive decoupling, evident in the appearance of tractions also in the inner lamella (i.e. closest to the cell-cell interface). In addition to fibers connecting FAs in the outer lamella to AJs in the cell-cell interface, there are fibers connecting FAs in the outer lamella to FAs in the inner lamella. The magnitude of traction exerted below the inner lamella become comparable to those in the outer lamella and are thus noticeable in the presented traction maps. This matches observations made on cell pairs in vitro (6, 7).

**Figure 3.**
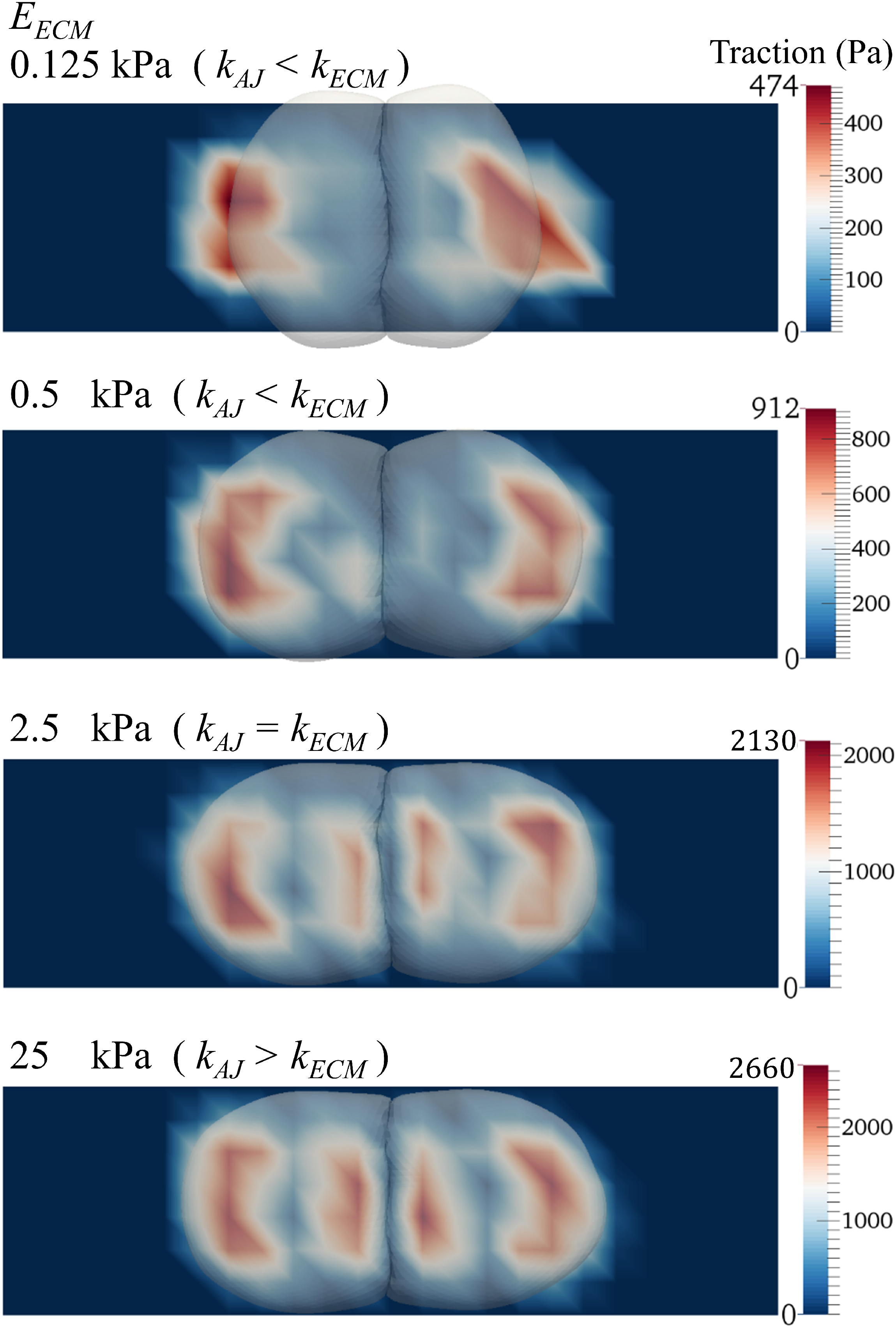
Traction maps for simulation of cell pairs on substrates of varying stiffness. Decoupling with increasing substrate stiffness (E_ECM_) is evidenced by the spatial redistribution of cellular tractions in the cell-substrate interface from underneath the outer region of the cell pair to underneath the cell-cell interface. These sample cases correspond to simulations with maturing FAs but not AJs (ζ_FA_ > ζ_AJ_). An animation of the entire simulation for each of these cases can be found in Supporting Material.

In all scenarios there was a qualitative agreement with experiments: By simulating cells pairs on increasingly stiff substrates the model resulted in decoupling of cells, increase in number of FAs, and increase in contractile strength of the cell pair.

To quantify this behavior, the results were analyzed using the same metrics used to characterize the behavior of cell pairs in vitro (6): intercellular force (*F*_*j*,*cell*_), the contractile moment of the cell pair (*M*_*xx*_), number of FAs, and cell coupling (*ψ*). The intercellular force (*F*_*j*,*cell*_) exerted by each cell was calculated as the unbalanced traction force exerted by the cell on the substrate through the total number of FAs (*n*_*FA*,*cell*_) of the cell, as described by Equation 10:

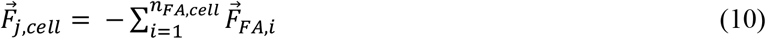

The contractile moment of the cell pair (*M*_*xx*_) provides a measurement of the total force exerted by the cell pair along its long axis, while taking into account the distance at which the traction force is exerted from the cell-cell interface (*r*_*x FA*,*i*_). Including this distance ensures deformation of the cell pair is taken into account in the comparisons between conditions. The contractile moment was calculated according to Equation 11:

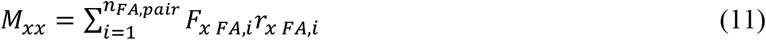

where 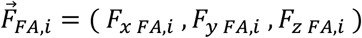. The distance *r*_*x FA*,*i*_ is signed and calculated along the *x*-axis (aligned with the long side of the rectangular pattern), and it is the distance from the average *x*-position of the cell-cell interface and the FA. More details on the calculation of *M*_*xx*_ can be found in the Supporting Material (Figure S4).

Because cells are known to exert higher tractions on stiffer substrates, the measure of intercellular force does not suffice to describe the coupling strength of cells. To account for this, a dimensionless ratio dubbed cell coupling (*ψ*) is used. This metric, defined in Equation 12, is the average over the two cells of the ratio of the magnitude of the intercellular force exerted by each cell 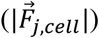 to the contractile moment of the cell normalized by cell length (*N*_*xx*,*cell*_). The normalized moment of each cell is defined in Equation 13. A higher cell coupling value (*ψ*) means relatively more force is exerted by the cells on each other; in contrast, a lower value of *ψ* means the cells are less coupled (relatively decoupled) with more force exerted by the cells on the substrate.

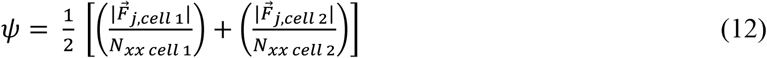

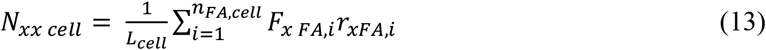

where *L*_*cell*_ is the length of the cell along the along the *x*-axis (Figure S4).

Values for intercellular force (*F*_*j*,*cell*_), contractile moment of the cell pair (*M*_*xx*_), number of FAs, and cell coupling (*ψ*) are displayed in Figure 4 (A-D). Reported values were obtained by averaging the results of the 5 replicates run for each condition: For each simulation, the values at every minute over the last hour were averaged to obtain a single one; this was done to account for fluctuations in the active system.

**Figure 4.**
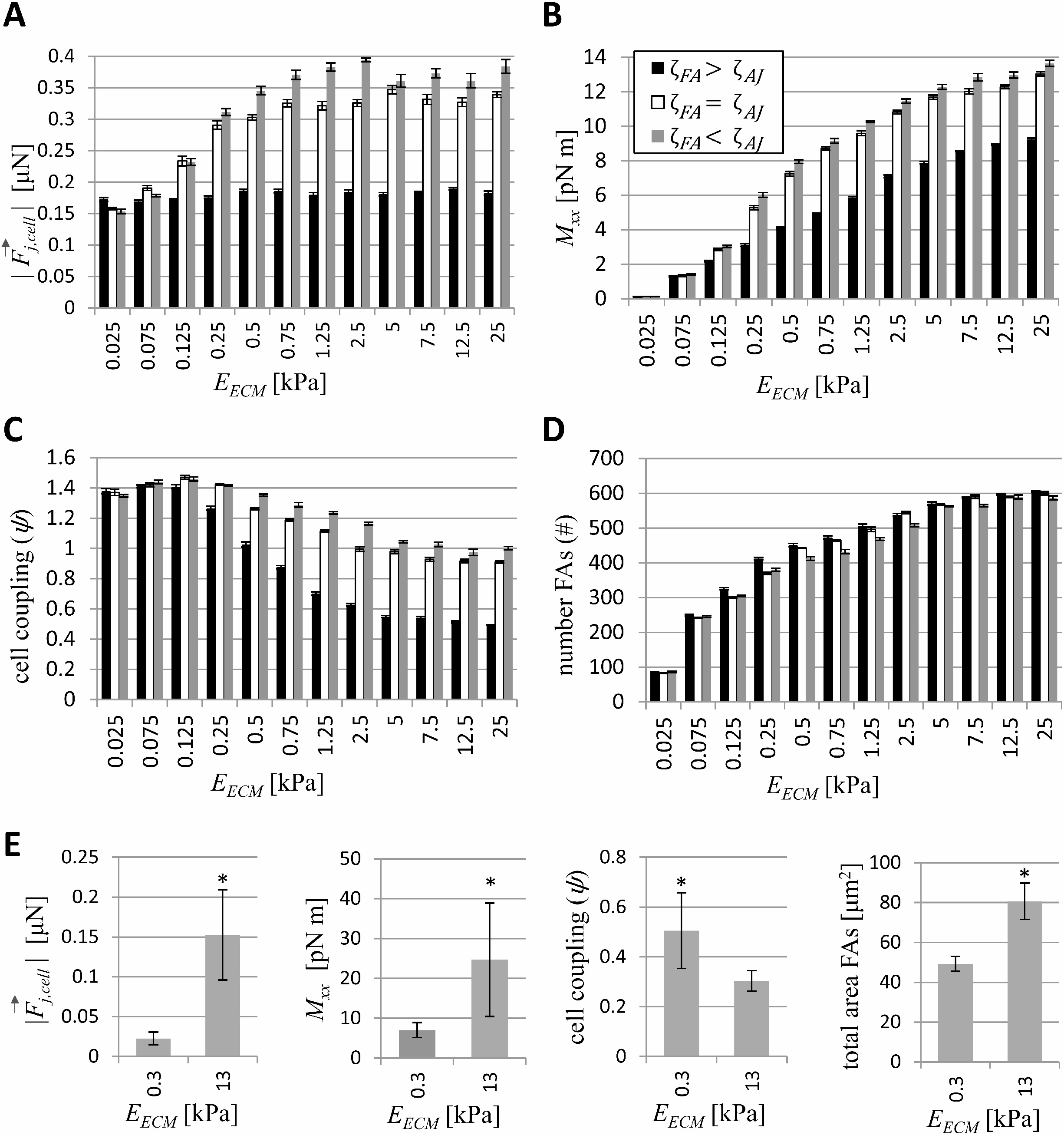
The mechanosensing capability of adherens junctions relative to focal adhesions was varied by varying the parameter ζ_AJ_, while keeping ζ_FA_ the same. Average values for simulated cell pairs of: A) Intercellular force (F_j,cell_), B) contractile moment of the cell pair (M_xx_), C) cell coupling (ψ), and D) number of focal adhesions bound to a stress fiber. n=5. Error bars correspond to standard error of the mean (SEM). E) Corresponding experimental results. Adapted from Polio et al. (6).

### Force dependent adhesion maturation regulates stress fiber dynamics

The results displayed in Figure 4 (A-C), show a drastic effect when making AJs mechanosing (ζ_FA_ ≤ ζ_AJ_) relative to when they are not (ζ_FA_ > ζ_AJ_). To explore what differs between these conditions, the stress fibers were more carefully analyzed. This quickly suggested that the connections between stress fibers and adhesions made a difference in terms of force response. Three different configurations in terms of connections can be defined: a) a stress fiber connecting two FAs (FA-FA) b) stress fibers that connect a FA to an AJ itself bound to another stress fiber (FA-AJ-FA), and c) stress fibers that connect a FA to an AJ not bound to any other fiber (FA-AJ). A graphical representation of these configurations can be found in the Supporting Material (Figure S5).

The factor describing fiber strengthening (*n*_*str*,*f*_) due to stalling during shortening was recorded and analyzed. For fibers in each configuration, there was a difference in this strengthening factor. These results for the first two configurations are presented in Figure 5 (A,B). In the case of the third configuration (i.e. fibers bound to an AJ only bound to a single fiber), no fiber matured (not shown). Also shown are the number of stress fibers in the cell pairs regardless of configuration in Figure 5C, and the number of fibers that connect a FA to an AJ itself bound to another stress fiber (FA-AJ-FA) in Figure 5D.

**Figure 5.**
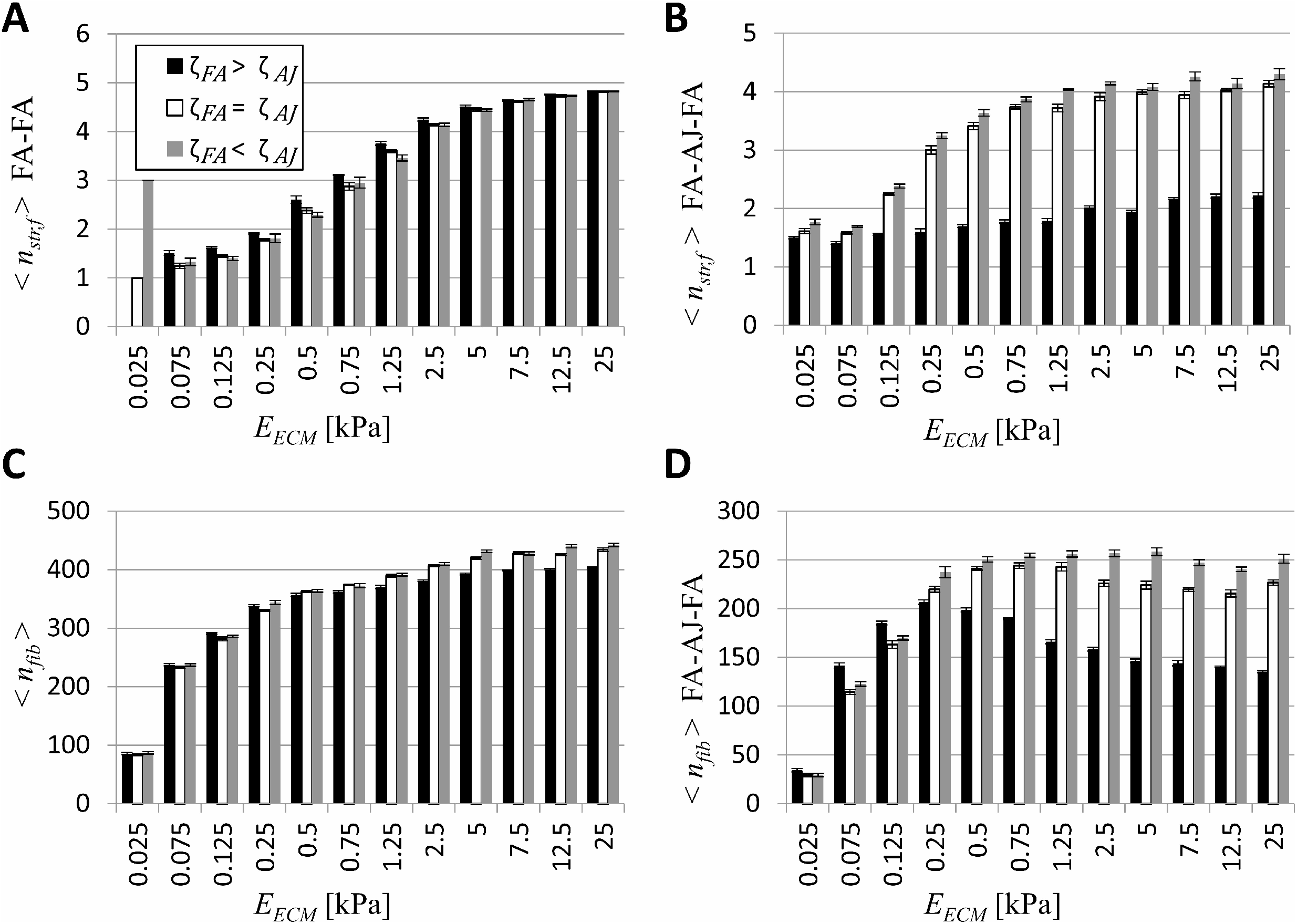
A) Average value of factor describing stress fiber strengthening for fibers in cell pair connecting two focal adhesions (< n_str,f_ > FA − FA). B) Average value of factor describing stress fiber strengthening for fibers in cell pair connected to another via an adherens junction (< n_str,f_ > FA − AJ − FA). C) Number of fibers in cell pair in all configurations (< n_fib_ >). D) Number of fibers in cell pair connected to another via an adherens junction (<n_fib_ > FA − AJ − FA). n=5. Error bars correspond to standard error of the mean (SEM).

The increase in number of stress fibers with increasing substrate stiffness (Figure 5C) can be attributed to the increase in fibers that connect a FA to an AJ itself bound to another stress fiber (FA-AJ-FA) on low stiffness substrates (Figure 5D) and to the increase in fibers connecting two FAs (FA-FA) on high stiffness substrates (Supporting Material, Figure S6A). Meanwhile, the number of fibers that connect a FA to an AJ not bound to any other fiber (FA-AJ) are few in comparison (Supporting Material, Figure S6B), as less resistance to contraction prevent the adhesions from maturing.

This change in the number of fibers in different configurations at different stiffness ranges shows that there is a competition for the limited number of nodes in the lamella to which stress fibers can bind once a FA is formed. On low stiffness substrates, stress fibers binding AJs in the cell-cell interface dominate cell pair contraction; meanwhile, as the substrate stiffness increases (and the stiffness sensed by fibers at the cell-cell interface remains the same), stress fibers binding two FAs (FA-FA) win. At this point, forces exerted on the substrate account for most of cellular contraction. This phenomenon is strongest when AJs are not stabilized by force (ζ_FA_ > ζ_AJ_), such that fibers bound to the cell-cell interface have less time to strengthen. Competition among the different fiber configurations can explain how simulated results match experimental observations.

### Stabilization of adherence junctions is needed for intercellular force increase

In Figure 4A the intercellular force (*F*_*j*,*cell*_) increases with substrate stiffness in the scenarios in which AJs matured with force (ζ_FA_ = ζ_AJ_ and ζ_FA_ < ζ_AJ_), but not in the scenario in which only FAs did (ζ_FA_ > ζ_AJ_). Meanwhile, Figure 5 (A,B) shows that there is an increase in strengthening of fibers with increasing substrate stiffness. Albeit slight, the increase in < *n*_*str*,*f*_ > FA-AJ-FA exists even in the scenario in which only FAs matured (ζ_FA_ > ζ_AJ_). This demonstrates that an average expected lifetime of AJs of ~17 min (corresponding to 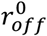 value in Table1 and comparable to the lifetime of adhesions biologically (25)) is in fact long enough for fibers to stall and strengthen.

When comparing between the scenarios in which AJs matured (ζ_FA_ = ζ_AJ_ and ζ_FA_ < ζ_AJ_), intercellular force was higher as the factor describing dynamics of mechanosensing (i.e. ζ_AJ_) increased; this occurred particularly at higher substrate stiffness values (*E*_*ECM*_ ≥ 0.25 kPa) and can be seen in Figure 4A. Intercellular force is determined by force exerted by fibers bound to the cell-cell interface (the majority in configuration FA-AJ-FA), so to understand how it varies we looked to maturation of these fibers. We did not necessarily expect ζ_*AJ*_ to have an effect on intercellular force, because force is exerted by the stress fibers which strengthen independently of the maturation of adhesions (i.e. *n*_*str*,*f*_ is independent of ζ_*AJ*_).

In both scenarios (ζ_FA_ = ζ_AJ_ and ζ_FA_ < ζ_AJ_), the factor *n*_*str*,*f*_ appears to plateau with increasing substrate stiffness (Figure 5B). The plateau in fibers in the configuration FA-AJ-FA is reached at a lower stiffness value relative to those in the configuration FA-FA: More specifically, in the former the plateau is reached at around 2.5 kPa, the value of the AJ spring constant (*k*_*AJ*_). The stiffness of the AJs (*k*_*AJ*_) acts as a sort of threshold above which stiffness of the substrate (*E*_*ECM*_) will no longer dictate intercellular force.

Experimentally, the intercellular force between cells in cell pairs increased from 0.022 to 0.15 μN (factor of ~6.7) when cells went from 0.3 to 13 kPa (Figure 4E). In simulations for the scenario in which FAs and AJs are equally mechanosensitive (ζ_FA_ = ζ_AJ_), there was an increase from 0.29 to 0.33 μN when cells went from 0.25 to 12.5kPa; this amounts to a factor of ~1.14. Over the entire substrate stiffness range studied, intercellular force increased by a factor closer to ~2.2.

### Increased intercellular force and increased contractile moment of the cell pair occur together

In all scenarios of maturation of AJs (relative to FAs), the contractile moment of the cell pair (*M*_*xx*_) increases with substrate stiffness (Figure 4B). In the scenario in which only the FAs mature (ζ_FA_ > ζ_AJ_), the contractile moment increases (unlike intercellular force) because this quantity depends on all configurations of stress fibers and adhesions and not only the configuration FA-AJ-FA (as was the case for intercellular force). Fibers bound to two FAs will increasingly strengthen with increasing substrate stiffness until they have reached the maximum factor (*n*_*str*,*f*_ = 5). Fibers in the configuration FA-FA on average near this maximum value of *n*_*str*,*f*_ = 5 at a stiffness value of *E*_*ECM*_ = 25 kPa (Figure 5A). In the scenarios in which AJs matured (ζ_FA_ = ζ_AJ_ and ζ_FA_ < ζ_AJ_), the contractile moment increased more drastically with increasing substrate stiffness, due to the contribution of fibers binding the interface. This is evidenced in the corresponding increase in intercellular force relative to the scenario in which only the FAs mature, already discussed (Figure 4A).

Experimentally, the contractile moment increased from 7.03 to 24.65 pN·m (factor of ~3.05) when cells went from 0.3 to 13 kPa (Figure 4B). In simulations for the scenario in which FAs and AJs are equally mechanosensitive (ζ_FA_ = ζ_AJ_), there was an increase from 5.29 to 12.28 pN·m when cells went from 0.25 to 12.5kPa, which amounts to a factor of ~2.32.

### The stiffness of the cellular interface relative to the substrate arbitrates cell coupling

Cell coupling (*ψ*) is defined in terms of intercellular force (*F*_*j*,*cell*_) and contractile moment of the cells (*N*_*xx*,*cell*_) (a quantity related to *M*_*xx*_) according to Equation 12. On low substrate stiffness (*E*_*ECM*_ ≤ 0.125 kPa), there is a slight increase in cell coupling with increase in substrate stiffness, and maturation of AJs (ζ_AJ_) has no effect on coupling (Figure 4C). A more drastic effect, however, is observed on stiffer substrates (0.125 kPa < *E*_*ECM*_ < 2.5 kPa) for which coupling decreases visibly with increasing substrate stiffness. This decrease is less drastic as the degree of mechanosensing of AJs increases (i.e. as ζ_AJ_ increases).

The change in coupling with substrate stiffness is again relatively small on the stiffest substrates, specifically at *E*_*ECM*_ ≥ 2.5 kPa, the value of the AJ spring constant (*k*_*AJ*_). This is expected from the definition of cell coupling in terms of intercellular force and contractile moment and their dependence on substrate stiffness: The trend in cell coupling values is a result of the limit reached by intercellular force (Figure 4A) and the limit in maturation of stress fibers (Figure 5 (A,B)) at this substrate stiffness. Experimentally, cell coupling also decreased: It decreased from 0.51 to 0.3 (factor of ~0.6) when cells went from 0.3 to 13 kPa (Figure 4B). In simulations for the scenario in which FAs and AJs are equally mechanosensitive (ζ_FA_ = ζ_AJ_), there was a decrease from 1.42 to 0.92 when cells went from 0.25 to 12.5kPa; this amounts to a factor of ~0.65. These results suggest that the substrate stiffness values probed experimentally (i.e. 0.3 and 13 kPa) lie on different sides of the perceived cell-cell interface stiffness and thus cause a difference in cell coupling.

These results suggest that AJ maturation acts counter to cell decoupling, yet it is needed for intercellular force to increase with increasing substrate stiffness. Cellular decoupling with increasing substrate stiffness could be attributed to strengthening of stress fibers. Fiber strengthening for fibers bound to the cell-cell interface, however, will be attenuated when the substrate becomes as stiff or stiffer than the perceived cell-cell interface (here dictated by AJ stiffness or *k*_*AJ*_) due to competition between the stress fibers in the different configurations (Supporting Material, Figure S5). At his point the increase in substrate stiffness will not lead to further strengthening of fibers as the contraction dynamics of the stress fibers and adhesions configuration will be dictated by the softest link: Fibers bound to AJs will not mature further, while those connecting two FAs will. This gives them the advantage and over time will outlive and replace those exerting force on the cell-cell interface. The cells then become decoupled. This also explains why there is no strengthening of stress fibers that bind a FA to an AJ not bound to any other fiber (FA-AJ): These fibers feel only the cell-cell interface, which has a stiffness dictated by the stiffness of the cell cortex (*k*_*cortex*_) and a series of other parameters (e.g. *k*_*b*_, *W*_*cc*_).

### Increase in number of FAs close to cell-cell interface accompanies decline in cell coupling

Unlike cells in vitro, in which a FA only exists if reinforced, in our simulations FAs can form based on the rate of assembly (*r*_*on*,*FA*_) and continue to exist if stabilized by force, exerted mainly by stress fibers bound to the adhesion itself. For this reason to compare the number of FAs with experimental measurements, the number of FAs in the cell pair that are bound to a fiber were tallied (Figure 4D).

The number of FAs was much lower at the softest substrate stiffness; here the substrate stiffness is so low that the cell easily deforms it, and the stress fibers keep contracting and never stall, preventing the strengthening of fibers connecting two FAs. This meant no fibers in FA-FA configuration remained at the end of the simulations at the lowest stiffness condition (*E*_*ECM*_ = 0.025 kPa) (Figure 6SA in Supporting Material). The cells contracted extensively, creating a larger cell-cell interface, such that they did not meet the minimum length criterion for stress fibers to form (see Methods). In substrates of all other stiffness values a cell-substrate interface large enough for lamella to develop and fibers to form and strengthen was formed. The number of FAs increased in the interval *E*_*ECM*_ (0.075, 25) kPa. Experimentally, the extent of discrete cell-substrate adhesion was measured via the area of FAs measured from fluorescence imaging: It increased from 49.03 to 80.7 μm^2^ (factor of ~1.64) when cells went from 0.3 to 13 kPa (Figure 4E). In simulations for the scenario in which FAs and AJs are equally mechanosensitive (ζ_FA_ = ζ_AJ_), the increase in number of FAs when cells went from 0.25 to 12.5kPa was from 369.91 to 589.15 adhesions; this amounts to a the factor of ~1.59.

The increase in the number of FAs can be attributed to the shift in number of the different categories of stress fibers, particularly the triumph of stress fibers binding two FAs (FA-FA) over those connecting FAs to the cell-cell interface on stiffer substrates. FAs bound to these fibers will be stabilized by the increased force with fiber strengthening. This is especially true for the inner lamella, in which only stress fibers binding two FAs can bind and exert force. This explains how in these areas close to the cell-cell interface there is increased traction exertion with increasing substrate stiffness, the results shown qualitatively to match experimental findings in Figure 3.

This phenomenon in which FAs form at a distance from where the intercellular junction, and AJs form, has been observed not only with changes in substrates stiffness as a driver but brought along by substrate geometry. Tseng *et al.* placed mammary epithelial (MCF10A) cell-pairs on square (outline), [H]-shaped, or [hourglass]-shaped (outline) micropatterns, instead of a rectangular pattern (filled), and found that the cell-cell adhesions will be positioned over areas where no ligand is present (i.e. outside the pattern where no FAs can form) (26). Although the mechanisms behind this observation remain unknown, researchers were able to show that the exertion of cell-cell forces is involved in the spatial guidance of cell-cell adhesions away from the substrate. Cells exerted lower forces at cell-cell interface (and overall) when the interface was located away from the ligand, possibly in an effort to minimize intracellular tension. In our simulations we see intercellular tension reaches a limit despite increasing substrate stiffness (Figure 4A), while the contractile moment of the cell pair continues to increase (Figure 4B). The result is a decoupling of cells in which competitive binding due to differing response to substrate stiffness of FAs and AJs cause the FAs to replace AJs near the cell-cell interface. This suggests that in these different geometries used by Tseng *et al.*, there may be a replacement as well driven by competitive binding and mediated by differing mechanosensing attributes of FAs and AJs.

### Cell decoupling will only occur in a range of substrate stiffness values

The range of substrate stiffness values was chosen after preliminary simulations showed that the cell pair stalled in its force response at both ends of the range. At the lowest value (*E*_*ECM*_ = 0.025), cells contract creating an exceptionally large cell-cell interface and a small cell-substrate interface; very few fibers in the FA-FA configuration could form and those binding the outer lamella to the cell-cell interface barely strengthened due to the soft substrate. This explains the sharp transition observed in the number of fibers (< *n*_*fib*_ >) observed on substrates between *E*_*ECM*_ = 0.025 and 0.075 kPa (Figure 5C). Comparing simulations on these same stiffness values in terms of strengthening of stress fibers connected via AJs, there is no difference in terms of maturation. Together, these observations suggest that no different behavior is expected at lower stiffness values than those selected in our simulations. At the high end of the range of substrate stiffness, no further change in behavior is expected either: All quantities presented in Figures 4 (A-D) and Figure 5 plateau. For this reason we believe the model is thorough in its exploration of the effect of stiffness on cell pair force exertion as a function of substrate stiffness.

## Conclusions

In this work we focused on the effect of mechanosensing on cell pairs, and specifically active mechanisms (i.e. adhesion maturation and stress fiber strengthening). The mechanical response of the cell, however, will be affected by additional elements, many passive. Our results show that stiffness at the cell-cell interface regulates the force response of the cell; in our simulations this stiffness was dictated by the stiffness of AJs. Future studies with the model may ask for the effect of some of these passive mechanical factors, such as cortex stiffness (*k*_*cortex*_). We wonder, could a varying cortex stiffness explain why different cell types show different behaviors on the same substrate stiffness? Additionally, the model can be expanded to collectives beyond cell pairs. The same way the specific configuration of stress fibers and adhesions are shown to play a role in force response to substrate stiffness, similar results can be expected of more complex configurations of these elements.

The model suggests that stalling and strengthening of stress fibers occurs faster in stress fibers that connect two FAs compared to in those binding AJs. This occurs because the cell-cell interface is softer than the substrate, but also because not all stress fibers bound to an AJ stall and mature; maturation requires stalling which requires an opposing force, and this only occurs when the AJ is bound to yet a second stress fiber in the neighbor cell. As this demands binding of two instead of one stress fiber, it is less likely thus causing stabilization of cell-cell adhesions to take longer. We also found limits in terms of substrate stiffness, relative to adhesion complex stiffness values, for which this occurs: The stiffness of the cell-cell interface, sensed as having a relatively constant stiffness, becomes a tipping point for substrate stiffness to define mechanosensing dynamics and cell strengthening. We do not expect these mechanisms to fully explain the complex interplay of cell-cell and cell-substrate adhesions; it is known that this interplay also requires chemical signaling and activation of common cellular pathways, most likely including RhoA kinase and Focal Adhesion Kinase pathways (8, 10, 27–29). Experimental measurements at the subcellular scale can be technically difficult to obtain. This model helps quantify the role of mechanosensing in the interplay of cell-cell and cell-ECM adhesions.

## Author Contributions

DAV, HP, and HVO designed the study. DAV, TH, and BS conceived the model. DAV implemented the model, ran simulations, and analyzed results. The group headed by HR developed the computational program used for coding the model and helped with trouble shooting.

## Acknowledgments

Funding for this work comes from the European Research Council (FP7/2007-2013 / ERC Grant Agreement n° 308223), FWO-Vlaanderen (Grant n° G.0821.13), and FWO and EU’s Horizon 2020 research and innovation programme (Marie Skłodowska-Curie Grant Agreement n° 665501). The computational resources and services used in this work were provided by the VSC (Flemish Supercomputer Center), funded by the Research Foundation - Flanders (FWO) and the Flemish Government – department EWI.

## References

1. Mui, K.L., C.S. Chen, and R.K. Assoian. 2016. The mechanical regulation of integrin-cadherin crosstalk organizes cells, signaling and forces. J. Cell Sci.: 1–8.

2. Matsushita, T., M. Oyamada, K. Fujimoto, Y. Yasuda, S. Masuda, Y. Wada, T. Oka, and T. Takamatsu. 1999. Remodeling of Cell-Cell and Cell–Extracellular Matrix Interactions at the Border Zone of Rat Myocardial Infarcts. Circ. Res. 85: 1046–1055.

3. Araujo, B.B., M. Dolhnikoff, L.F.F. Silva, J. Elliot, J.H.N. Lindeman, D.S. Ferreira, A. Mulder, H.A.P. Gomes, S.M. Fernezlian, A. James, and T. Mauad. 2008. Extracellular matrix components and regulators in the airway smooth muscle in asthma. Eur. Respir. J. 32: 61–9.

4. An, S.S., W. Mitzner, W.-Y. Tang, K. Ahn, A.-R. Yoon, J. Huang, O. Kilic, H.M. Yong, J.W. Fahey, S. Kumar, S. Biswal, S.T. Holgate, R.A. Panettieri, J. Solway, and S.B. Liggett. 2016. An inflammation-independent contraction mechanophenotype of airway smooth muscle in asthma. J. Allergy Clin. Immunol. 138: 294–297.e4.

5. Tan, C., P. Costello, J. Sanghera, D. Dominguez, J. Baulida, A. Garcia de Herreros, and S. Dedhar. 2001. Inhibition of integrin linked kinase (ILK) suppresses β-catenin-Lef/Tcf-dependent transcription and expression of the E-cadherin repressor, snail, in APC−/− human colon carcinoma cells. Oncogene. 20: 133–140.

6. Polio, S.R., S.E. Stasiak, R.R. Jamieson, J.L. Balestrini, R. Krishnan, and H. Parameswaran. 2019. Extracellular matrix stiffness regulates human airway smooth muscle contraction by altering the cell-cell coupling. Sci. Rep. 9: 9564.

7. McCain, M.L., H. Lee, Y. Aratyn-Schaus, A.G. Kléber, and K.K. Parker. 2012. Cooperative coupling of cell-matrix and cell-cell adhesions in cardiac muscle. Proc. Natl. Acad. Sci. U. S. A. 109: 9881–6.

8. Krishnan, R., D.D. Klumpers, C.Y. Park, K. Rajendran, X. Trepat, J. van Bezu, V.W.M. van Hinsbergh, C. V. Carman, J.D. Brain, J.J. Fredberg, J.P. Butler, and G.P. van Nieuw Amerongen. 2011. Substrate stiffening promotes endothelial monolayer disruption through enhanced physical forces. Am. J. Physiol. Physiol. 300: C146–C154.

9. Goeckeler, Z.M., and R.B. Wysolmerski. 1995. Myosin light chain kinase-regulated endothelial cell contraction: the relationship between isometric tension, actin polymerization, and myosin phosphorylation. J. Cell Biol. 130: 613–27.

10. Kugelmann, D., L.T. Rotkopf, M.Y. Radeva, A. Garcia-Ponce, E. Walter, and J. Waschke. 2018. Histamine causes endothelial barrier disruption via Ca2+-mediated RhoA activation and tension at adherens junctions. Sci. Rep. 8: 13229.

11. Yonemura, S., Y. Wada, T. Watanabe, A. Nagafuchi, and M. Shibata. 2010. α-Catenin as a tension transducer that induces adherens junction development. Nat. Cell Biol. 12: 533–542.

12. Odenthal, T., B. Smeets, P. Van Liedekerke, E. Tijskens, H. Van Oosterwyck, and H. Ramon. 2013. Analysis of initial cell spreading using mechanistic contact formulations for a deformable cell model. PLoS Comput. Biol. 9: e1003267.

13. Delorme, V., M. Machacek, C. DerMardirossian, K.L. Anderson, T. Wittmann, D. Hanein, C. Waterman-Storer, G. Danuser, and G.M. Bokoch. 2007. Cofilin Activity Downstream of Pak1 Regulates Cell Protrusion Efficiency by Organizing Lamellipodium and Lamella Actin Networks. Dev. Cell. 13: 646–662.

14. Van Liedekerke, P., E. Tijskens, H. Ramon, P. Ghysels, G. Samaey, and D. Roose. 2010. Particle-based model to simulate the micromechanics of biological cells. Phys. Rev. E. 81: 061906.

15. Maugis, D. 1992. Adhesion of spheres: The JKR-DMT transition using a dugdale model. J. Colloid Interface Sci. 150: 243–269.

16. Schwarz, U.S., T. Erdmann, and I.B. Bischofs. 2006. Focal adhesions as mechanosensors: The two-spring model. Biosystems. 83: 225–232.

17. Mitrossilis, D., J. Fouchard, A. Guiroy, N. Desprat, N. Rodriguez, B. Fabry, and A. Asnacios. 2009. Single-cell response to stiffness exhibits muscle-like behavior. Proc. Natl. Acad. Sci. U. S. A. 106: 18243–8.

18. Seddiki, R., G.H.N.S. Narayana, P.-O. Strale, H.E. Balcioglu, G. Peyret, M. Yao, A.P. Le, C. Teck Lim, J. Yan, B. Ladoux, and R.M. Mège. 2018. Force-dependent binding of vinculin to α-catenin regulates cell–cell contact stability and collective cell behavior. Mol. Biol. Cell. 29: 380–388.

19. Elosegui-Artola, A., R. Oria, Y. Chen, A. Kosmalska, C. Pérez-González, N. Castro, C. Zhu, X. Trepat, and P. Roca-Cusachs. 2016. Mechanical regulation of a molecular clutch defines force transmission and transduction in response to matrix rigidity. Nat. Cell Biol. 18: 540–548.

20. Bangasser, B.L., S.S. Rosenfeld, and D.J. Odde. 2013. Determinants of Maximal Force Transmission in a Motor-Clutch Model of Cell Traction in a Compliant Microenvironment. Biophys. J. 105: 581–592.

21. Wolfenson, H., G. Meacci, S. Liu, M.R. Stachowiak, T. Iskratsch, S. Ghassemi, P. Roca-Cusachs, B. O’Shaughnessy, J. Hone, and M.P. Sheetz. 2016. Tropomyosin controls sarcomere-like contractions for rigidity sensing and suppressing growth on soft matrices. Nat. Cell Biol. 18: 33–42.

22. Parameswaran, H., K.R. Lutchen, and B. Suki. 2014. A computational model of the response of adherent cells to stretch and changes in substrate stiffness. J. Appl. Physiol. 116: 825–34.

23. Purcell, E.M. 1977. Life at low Reynolds number. Am. J. Phys. 45: 3–11.

24. Austen, K., P. Ringer, A. Mehlich, A. Chrostek-Grashoff, C. Kluger, C. Klingner, B. Sabass, R. Zent, M. Rief, and C. Grashoff. 2015. Extracellular rigidity sensing by talin isoform-specific mechanical linkages. Nat. Cell Biol. 17: 1597–1606.

25. Webb, D.J., K. Donais, L.A. Whitmore, S.M. Thomas, C.E. Turner, J.T. Parsons, and A.F. Horwitz. 2004. FAK–Src signalling through paxillin, ERK and MLCK regulates adhesion disassembly. Nat. Cell Biol. 6: 154–161.

26. Tseng, Q., E. Duchemin-Pelletier, A. Deshiere, M. Balland, H. Guillou, O. Filhol, and M. Théry. 2012. Spatial organization of the extracellular matrix regulates cell-cell junction positioning. Proc. Natl. Acad. Sci. U. S. A. 109: 1506–11.

27. Mikelis, C.M., M. Simaan, K. Ando, S. Fukuhara, A. Sakurai, P. Amornphimoltham, A. Masedunskas, R. Weigert, T. Chavakis, R.H. Adams, S. Offermanns, N. Mochizuki, Y. Zheng, and J.S. Gutkind. 2015. RhoA and ROCK mediate histamine-induced vascular leakage and anaphylactic shock. Nat. Commun. 6: 6725.

28. Chen, X.L., J.-O. Nam, C. Jean, C. Lawson, C.T. Walsh, E. Goka, S.-T. Lim, A. Tomar, I. Tancioni, S. Uryu, J.-L. Guan, L.M. Acevedo, S.M. Weis, D.A. Cheresh, and D.D. Schlaepfer. 2012. VEGF-Induced Vascular Permeability Is Mediated by FAK. Dev. Cell. 22: 146–157.

29. Tseng, Q., E. Duchemin-Pelletier, A. Deshiere, M. Balland, H. Guillou, O. Filhol, and M. Théry. 2012. Spatial organization of the extracellular matrix regulates cell-cell junction positioning. Proc. Natl. Acad. Sci. U. S. A. 109: 1506–11.

30. Pontes, B., P. Monzo, and N.C. Gauthier. 2017. Membrane tension: A challenging but universal physical parameter in cell biology. Semin. Cell Dev. Biol. 71: 30–41.

31. Fischer, R.S., M. Gardel, X. Ma, R.S. Adelstein, and C.M. Waterman. 2009. Local Cortical Tension by Myosin II Guides 3D Endothelial Cell Branching..

32. Dembo, M., and Y.-L. Wang. 1999. Stresses at the Cell-to-Substrate Interface during Locomotion of Fibroblasts. Biophys. J. 76: 2307–2316.

33. Moore, S.W., P. Roca-Cusachs, and M.P. Sheetz. 2010. Stretchy Proteins on Stretchy Substrates: The Important Elements of Integrin-Mediated Rigidity Sensing. Dev. Cell. 19: 194–206.

